# The 2-approximation algorithm of sorting by prefix transposition revisited

**DOI:** 10.1101/036814

**Authors:** Md. Shafiqul Islam, Md. Khaledur Rahman, M. Sohel Rahman

## Abstract

A transposition is an operation that exchanges two adjacent blocks in a permutation. A prefix transposition always moves a prefix of the permutation to another location. In this article, we use a data structure, called the permutation tree, to improve the running time of the best known approximation algorithm (with approximation ratio 2) for sorting a permutation by prefix transpositions. By using the permutation tree, we improve the running time of the 2-approximation algorithm to *O*(*n* log *n*).

## I. Introduction

One of the most promising ways to trace evolutionary events is to compare the order of appearance of identical genes in two different genomes. In the 1980s, evidence was found that different species have essentially the same set of genes, but their order may differ between species [Palm er and Herbon 1986; Hoot and Palmer 1994]. This suggests that global rearrangement events such as *reversal, transposition*, and *block interchange* can be used to trace the evolutionary path between genomes. Such rare events may provide more accurate clues to evolution than local mutations (i.e., *insertions, deletions*, and *substitutions* of nucleotides).

In the last decade, a large body of work was devoted to genome rearrangement problems. The basic task here is as follows. Given two permutations, find a shortest sequence of rearrangement operations that transforms one given permutation into the other. Assuming that one of the permutations is the identity permutation, the problem is to find the shortest way of sorting a permutation using specific given rearrangement operations.

The best studied rearrangement event is the reversal. A reversal inverts a block of any size in a genome. Caprara proved that finding the minimum number of reversals needed to transform one genome into another is an NP-Hard problem [8]. Bafna and Pevzner presented an algorithm with approximation ratio 2 for this problem [9]. Later Berman, Hannenhalli and Karpinski presented the best known algorithm for the problem, with approximation ratio 1.375 [10].

Another interesting variation of the problem involves applying prefix reversal as the gnome rearrangement operator. In this problem, also known as the pancake flipping problem in the literature, only reversals involving the first consecutive elements of a genome are permitted. Gates and Papadimitriou [12] and Heydari and Sudborough [13] have studied the diameter of prefix reversals.

Another interesting and much studied rearrangement event is transposition. Transposition refers to the event of exchanging two adjacent blocks of any size in a genome. The transposition distance problem, that is, the problem of finding the minimum number of transpositions necessary to transform one genome into another, has been studied by Bafna and Pevzner [5]. After a long-standing 15-years of being unclassified this problem has very recently been classified to be NP-Hard in [15]. The best known algorithm for sorting a permutation by transposition has an approximation ratio 1.375 with running time *O*(*n* log *n*) [3] [14].

Like the prefix reversal, prefix transposition has also been given some attention in the literature. Dias, Fortuna and Meidanis presented an algorithm to sort a permutation by prefix transposition having approximation ratio 2 [2]. They also presented an algorithm to sort the reverse permutation, *R_n_* = [*n*, *n*–1,……., 3, 2,1] with 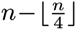 prefix transpositions.

In the literature, the problems of sorting by different rearrangement operations have been tackled from at least to different directions. In one direction, researchers have been trying to improve the approximation ratio and in the other, efforts have been made to improve the running time of the algorithm keeping the approximation ratio intact. In this paper, we are concerned with the latter direction. In [2], the authors did not formally present and analyze their 2-approximation algorithm for the problem of sorting by prefix transposition. If we implement this algorithm without any sophisticated data structure, the running time becomes *O*(*n*^2^) in the worst case. In this paper, we use an efficient data structure called permutation tree to sort a permutation by prefix transpositions. The motivation of our approach comes from a recent work of Firoz et al [3] where permutation tree has been used to achieve a faster running time for the best known approximation algorithm for sorting by transpositions. Using the permutation tree, in this paper, we present an *O*(*n* log *n*) time implementation of the 2-approximation algorithm of [2] for sorting by prefix transposition.

The rest of the paper is organized as follows. In Section II, we define important concepts that will be used throughout the paper. In Section III, we briefly review the permutation tree data structure. In Section IV, we give several results for 2-approximation algorithm of [2]. Finally, we briefly conclude in Section VI.

## II. Preliminaries

In this section, we introduce and define some notions and definitions that will be used throughout the paper. An arbitrary genome formed by n genes will be represented as a permutation *π* = [*π*(1),*π*(2),…,*π*(*n*)] where each element of *π* represents a gene. The identity genome *t_n_* is defined as *t_n_* = [1, 2, 3, …*n*]. A transposition *τ*(*x*,*y*,*z*), where 1 ≤ *x* ≤ *y* ≤ *z* ≤ *n* +1, is a rearrangement event that transforms *π* into the genome *πτ* = [*π*(1),…, *π*(*x* – 1),*π*(*y*),…,*π*(*z* – 1),*π*(*x*,…*π*(*y* – 1),*π*(*z*,…,*π*(*n*)]. In case of prefix transposition, *x* = 1 in *τ* (*x*, *y*, *z*). Given two genomes *π* and *σ* we define the prefix transposition distance *d_p_*(*π*,*σ*) between these two genomes as being the least number of prefix transpositions needed to transform *π* into *σ*, that is, the smallest *r* such that there are prefix transpositions *τ*_1_,*τ*_2_,…,*τ_r_* with *τ_r_*…*τ*_2_*τ*_1_*π* = *σ*. We call sorting distance by prefix transpositions, *d_p_*(*π*), the prefix transposition distance between the genomes *π* and *t_n_*, that is, *d_p_*(*π*) = *d_p_*(*π*, *t_n_*).

A breakpoint for the prefix transposition problem is a position *i* of a permutation *π* such that *π*(*i*) – *π*(*i* – 1) ≠ 1, where 2 ≤ *i* ≤ *n*. By definition, position 1 (beginning of the permutation) is always considered a breakpoint. Position *n*+1 (end of the permutation) is considered a breakpoint when *π*(*n*) = *n*. We denote by *b_p_*(*π*) the number of breakpoints of a permutation *π*. By definition *b_p_*(*π*) ≥ 1 for any permutation *π* and the only permutations with exactly one breakpoint are the identity permutations (*t_n_*, for all *n* ≥ 1). It is easy to see that there are no permutations with exactly two breakpoints. A strip is a subsequence *π*(*i*…*j*) of *π* where *i* ≤ *j* such that *i* and *j* + 1 are breakpoints and there are no breakpoints between positions *i* and *j*. Given a permutation *π* and a prefix transposition *τ*, we use Δ*b_p_*(*π*, *τ*) to denote the change on the number of breakpoints due to operation *τ*, that is, Δ*b_i_*(*π*,*τ*) = *b_p_*(*τπ*) – *b_p_*(*π*).

## III. Permutation Tree

A permutation tree is firstly a balanced binary tree *T* with root *r*, where each internal node of *T* has two children. Let t be a node of *T*. The left and right children of *t* are denoted as *L*(*t*) and *R*(*t*), respectively. The height of a leaf node is defined to be zero. The height of an internal node *t*, denoted *H*(*t*), is calculated as follows: *H*(*t*) = max (*H*(*L*(*t*)), *H*(*R*(*t*)) +1. Moreover, the tree must have the property of balance, that is, for any node *t* of *T*, |*H*(*L*(*t*)) – *H*(*R*(*t*))| ≤ 1. The height of *T* is defined to be the height of the root, that is, *H*(*T*) = *H*(*r*).

Secondly, a permutation tree must correspond to a permutation. For the permutation *π* = [*π*(1),*π*(2),…,*π*(*n*)], the permutation tree corresponding to *π* has *n* leaf nodes, which are labeled by *π*(1),*π*(2),…,*π*(*n*), respectively. Each node of *T* corresponds to an interval of *π* and is labeled by the maximum number in the interval. For any internal node *t* of *T*, the interval corresponding to *t* must be the concatenation of the two intervals corresponding to *L*(*t*) and *R*(*t*). The number labeled to *t* is called the value of *t*. Clearly, the value of node *t* must be the maximum value of *L*(*t*) and *R*(*t*). In the following, we may directly use the element *π*(*i*) to represent the leaf node labeled by *π*(*i*) and use the node *t* of *T* to represent the subtree of *T* rooted at *t*. As an example, Figure 1 is a permutation tree corresponding to the permutation *π* = [9, 6,1, 4, 7, 5, 2, 3, 8].

**Fig. 1.**
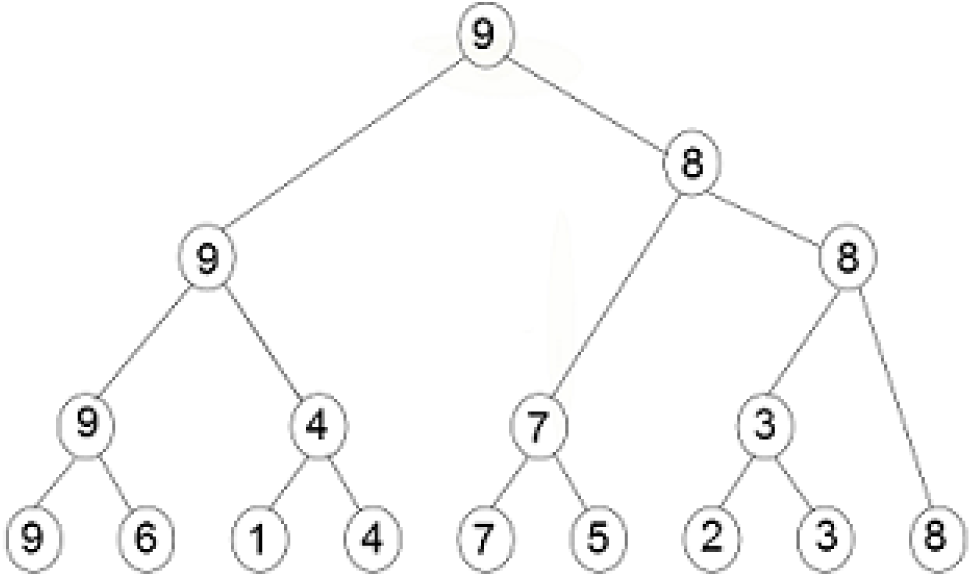
A permutation tree for *π* = [9; 6; 1; 4; 7; 5; 2; 3; 8].

The distinctive property of the permutation tree is that one tree represents a permutation, and every node of the tree represents an interval of the permutation. In addition, we do not need to insert or delete keys in the permutation tree. Feng and Zhu introduced three operations for a permutation tree [1]. These are *Build*, which builds a permutation tree corresponding to a given permutation, *Join*, which joins two trees into one, and *Split*, which splits one tree into two. The following results on permutation tree will be used later.

**Theorem III.1** ([1]). *The height of the permutation tree corresponding to π* = [*π*(1),*π*(2),…,*π*(*n*] *is bounded by O*(log *n*).

**Theorem III.2** ([1]). *Build always creates a permutation tree corresponding to a given permutation of* [1,…,*n*] *in O*(*n*) *time*.

**Theorem III.3** ([1]). *If t*_1_ *corresponds to* 1,…,*m*, *and t*_2_ *corresponds to m* + 1, …,*n*, *then Join*(*t*_1_,*t*_2_) *returns a permutation tree corresponding to* 1, …, *m*, *m* + 1, …, *n in O*(*H*(*t*_1_) – *H*(*t*_2_)) *time*.

**Theorem III.4** ([1]). *Suppose T is the permutation tree corresponding to ρ* = [*π*(1),…, *π*(*m* – 1), *π*(*m*),…, *π*(*n*)]. *The function Split*(*T*, *m*) *always returns T_l_ corresponding to ρ_ℓ_* = [*π*(1),…, *π*(*m* – 1)], *and T_r_ corresponding to ρ_r_* = [*π*(*m*),…,*π*(*n*)] *in O*(log *n*) *time*.

## IV. 2-Approximation Algorithm of [2]

The following results are due to [2] and lay the foundation for the 2-approximation algorithm for sorting by prefix transposition.
**Lemma IV.1** ([2]). *Given a permutation π* ≈; *t_n_*, *where n* = |*π*|, *it is always possible to obtain a prefix transposition τ such that* Δ*b_p_*(*π*,*τ*) ≤ –1.

**Lemma IV.2** ([2]). *Let π be a permutation with b_p_*(*π*) = 3, *then d_p_*(*π*) = 1.

**Lemma IV.3** ([2]). *For every permutation π different from the identity permutation we have d_p_*(*π*) ≤ *b_p_*(*π*) – 2.

**Lemma IV.4** ([2]). *For every permutation π different from the identity permutation we have 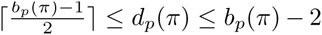*

**Theorem IV.5** ([2]). *Any algorithm that produces the prefix transpositions according to Lemmas IV. 1 and IV. 2 is an approximation algorithm with factor* 2 *for the prefix transposition distance problem*.

## V. Implementation

In this section, we discuss possible implementations of the algorithm of [2]. We first discuss the implementation without any sophisticated data structure and deduce the worst case time complexity of the implementation in Section V-A. Then, in Section V-B we show how permutation tree can be employed to get a time complexity of *O*(*n* log *n*) for the same algorithm.

### A. Implementation Without Permutation Tree

We use an 1-D array to hold the permutation and perform prefix transpositions on that array. For each prefix transposition, first we find the last element of the first strip as *k* and then, find *i* and *j*, where *i* = *π*^-1^(*k*) + 1 and *j* = *π*^-1^(*k* + 1). Then, interchange block [*π*(1),…,*π*(*π* – 1)] and [*π*(*i*),…, *π*(*i* – 1)] by using a simple *loop*. This *loop* takes *O*(*n*) time. According to *Lemma IV*. 1, each such prefix transposition reduces the number of break points from the permutation by at least one. So, total number of prefix transpositions needed is at most *n*. As a result, time complexity of this approach is *O*(*n*^2^).

### B. Implementation With Permutation Tree

In *Algorithm* 1 we present how to perform a prefix transposition operation using a permutation tree. In this algorithm, the procedure *Split*(*T*, *i*) receives a permutation tree *T* and an integer *i*, and returns two permutation trees *T*1 and *T*2, that represent the permutations [*π*(1),*π*(2), …,*π*(*i* – 1)] and [*π*(*i*), *π*(*i* + 1),…*π*(*n*)] respectively; *Join*(*T*1, *T*2) receives two permutation trees *T*1, *T*2 representing permutations [*π*(1), *π*(2),…, *π*(*i* – 1)] and [*π*(*i*), *π*(*i*+1),…, *π*(*n*) respectively, and returns a permutation tree *T* that represents [*π*(1), *π*(2),…, *π* (*i* – 1), *π*(*i*),…, *π*(*n*)]. Then we present *Algorithm* 2 to perform the actual sorting of a permutation by prefix transpositions. The procedure *Build*(*π*) used in *Algorithm* 2 receives a permutation *π* and returns a permutation tree. *π*^-1^(*k*) denotes the index position of element *k* in the permutation *π*. This algorithm runs until *π* becomes a permutaion of sorted list i,e identity permutaton. In the rest of this section, we state and prove a number of *Lemmas* concerning the running time of *Algorithm* 1 and *Algorithm* 2, achieving an *O*(*n* log *n*) running time of the overall algorithm in the sequel.

**Algorithm 1.**
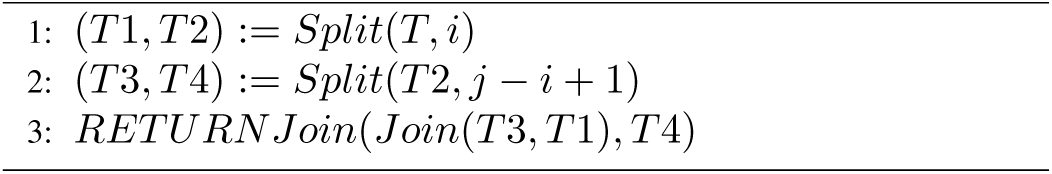
PREFIX-TRANSP0SITI0N(*T, i, j*)

**Algorithm 1.**
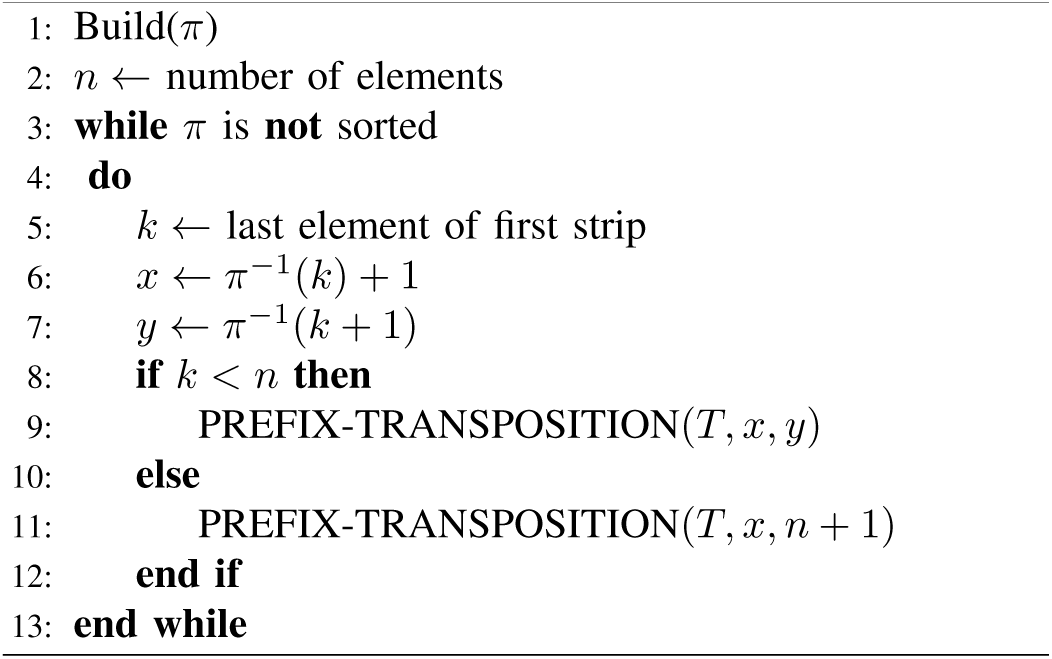
SORTING PERMUTATION BY PREFIX TRANSPOSITION

**Lemma V.1**. *Algorithm* 1 *can be implemented in O*(log *n*) *time*.

*Proof:* In *Algorithm* 1, we see that a prefix transposition takes two *Split* and two *Join* operations. From *Theorem III*.3, a *Join* operation can be implemented in *O*(*H*(*t*1) – *H*(*t*2))) time and from *Theorem III*.4, a *Split* operation can be implemented in *O*(log *n*) time. So time complexity is 2 × *O*(*H*(*t*1) – *H*(*t*2))) + 2 × *O*(log *n*), which is *O*(log *n*).

**Lemma V.2**. *Algorithm* 2 *can be implemented in O*(*n* log *n*) *time*.

*Proof:* The first step of Algorithm 2 is *Build*(*π*) operation, which takes *O*(*n*) time [1]. Steps 5 – 7 needs constant time to run. According to *Lemma V*.1 steps 8 – 11 can be implemented in *O*(log *n*) time. So each iteration of the *while* loop in Line 3 takes *O*(log *n*) time. According to *Lemma IV*.1, each prefix transposition reduces the number of break points from the permutation by at least one. So the total number of prefix transpositions needed is at most *n*. So, the loop iterates at most n times and since each iteration takes *O*(log *n*) time, the running time of the *while* loop is *O*(*n* log *n*). So the time complexity of *Algorithm* 2 is *O*(*n*) + *O*(*n* log *n*), which is *O*(*n* log *n*).

## VI. Conclusions

We introduced the time complexity of sorting a permutation by prefix transposition using permutation tree. Sorting a permutation by prefix transpositions, implemented with permutation tree, runs in *O*(*n* log *n*) time.

## ACKNOWLEDGMENT

We express our gratefulness and honor to Dr. M. Sohel Rahman, Associate Professor, Department of Computer Science and Engineering, Bangladesh University of Engineering and Technology, for his support, advice and care. His patience, scholarly guidance, encouragement, constructive criticism, valuable advice, reading inferior drafts and correcting them at all stages help us properly finish this work.

The Department of Computer Science and Engineering of Bangladesh University of Engineering and Technology provided an ideal environment for conducting our research. Faculty, students and staff alike have created an incredibly friendly and collaborative work environment.

